# BGC Atlas v2: biosynthetic gene clusters with taxonomic and environmental context at scale

**DOI:** 10.64898/2026.09.14.751540

**Authors:** Caner Bağcı, Alec Talamás-Tanner, Nadine Ziemert

## Abstract

Genomic and metagenomic studies have revealed that microbes have the capacity to produce an enormous diversity of secondary metabolites. These compounds play important roles in microbial interactions and are also a major source of medicines and other useful natural products, yet only a small fraction of this biosynthetic potential has been experimentally characterized. BGC Atlas was developed to explore this largely uncharacterized diversity by placing biosynthetic gene clusters (BGCs) into genomic and environmental context. Here, we present BGC Atlas v2, which expands the collection more than eightfold to more than 16 million predicted BGCs and extends it from metagenomic assemblies to metagenome-assembled genomes (MAGs), single-amplified genomes and isolate genomes. The new release provides taxonomic assignments for nearly all BGCs, harmonized environmental metadata, nested searches across biosynthetic, taxonomic, environmental and geographic properties, and protein-sequence searches against BGC-encoded genes. Its scale also reveals how much bacterial and archaeal biosynthetic diversity remains unexplored. By connecting pathways to related families, organisms and environments, BGC Atlas v2 supports natural-product discovery and investigations of prokaryotic biosynthetic diversity across taxa and ecosystems. BGC Atlas v2 is freely available at https://bgc-atlas.ziemertlab.com.

## 1 Introduction

Microbial secondary metabolites are fundamental components of microbial ecology, mediating interactions with competitors, hosts, and the environment. They are also important sources of pharmaceutical and biotechnological products [1, 2]. Their biosynthesis is generally encoded by biosynthetic gene clusters (BGCs), neighbouring sets of genes that encode the biosynthetic enzymes, regulators, and transporters required for the production of these metabolites. The organization of BGCs makes them identifiable in sequence data, and genome mining has become a central approach for investigating microbial secondary metabolism [3]. Metagenomics enables this approach to be extended beyond cultivated isolates, revealing extensive taxonomic and biosynthetic diversity across terrestrial, marine, and host-associated microbiomes [4–7]. Genome and metagenome sequencing have revealed an extraordinary diversity of biosynthetic pathways encoded by microbes, highlighting the scale of the chemical potential present in nature. However, identifying a BGC is only the first step towards understanding its biological and chemical significance. Interpreting a newly identified pathway requires information about whether related BGCs have been observed before, which organisms encode them, and in which environments they occur.

The first release of BGC Atlas [8] addressed this need through a centralized resource for exploring the diversity of BGCs in metagenomic datasets across different habitats. It identified 1,854,079 BGCs in 31,316 metagenome assemblies, organized them into 18,566 gene cluster families (GCFs), and linked them to their associated sample metadata. This enabled users to examine the environmental distributions of BGCs and their GCFs, inspect individual families, and query user-supplied BGCs against a large metagenomic collection. Since its release, BGC Atlas has been used as a reference collection for assessing BGC novelty, retrieving metagenomic homologs of BGCs of interest, and providing environmental, biogeographical, and taxonomic context to newly identified clusters [9–15]. Its underlying data have also been reused to augment training data for machine-learning-based annotation of BGC products [16]. At the same time, these applications, together with feedback from users, highlighted what the first release lacked: BGCs carried no taxonomic assignment and could not be attributed to an organism, the collection contained no genomes, and its coverage was restricted to the metagenome assemblies available in MGnify [17].

Here, we present BGC Atlas v2, which expands the collection more than eightfold to more than 16 million BGCs and incorporates more and larger metagenomic datasets, as well as metagenome-assembled genomes (MAGs), single-amplified genomes (SAGs), and isolate genomes. The new release adds taxonomic assignments to the identified BGCs, harmonizes environmental metadata across multiple data sources, and uses an updated GCF clustering approach to organize biosynthetic diversity at substantially increased scale. A redesigned web application provides additional search modes and expanded options for searching and filtering samples, genomes, studies, BGCs, and GCFs. Together, these developments allow users to place individual biosynthetic pathways into a broader genomic, taxonomic, and environmental context and to explore how microbial biosynthetic potential is distributed across environments and taxa.

## 2 Materials and methods

### 2.1 Data sources

We assembled the underlying sequence collection from environmental metagenome assemblies and microbial genome collections originating from different public sources. Environmental metagenome assemblies were obtained from MGnify [17] and from SPIRE assemblies [18] represented in the “Other environmental samples” subset of Metalog [19]. For MGnify-derived assemblies, sample metadata were retrieved through the MGnify application programming interface (API), whereas for SPIRE assemblies they were obtained from Metalog.

To broaden environmental and sequencing-type coverage, we additionally incorporated manually collected metagenome assemblies from the Microflora Danica project [20], global deep-sea hydrothermal deposits [21], the FRAM Strait time series [22], a high-resolution surface-ocean survey [23], and two Schönbuch forest soil datasets [9, 24], as well as long-read metagenomes from Great Barrier Reef seawater [25] and the 2020 Helgoland spring phytoplankton bloom [26]. For the FRAM Strait time series, associated environmental measurements were obtained from PANGAEA [27]. Collection-specific source identifiers and provenance were retained for all assemblies (Supplementary Table S1).

The genome collection included MAGs and isolate genomes from the high-quality subset of the Genome Taxonomy Database (GTDB) [28], the Ocean Microbiomics Database (OMDB) [6, 29], SPIRE [18], GEM [5], SMAG [30], TPMC [31], the Microflora Danica short-[20] and long-read MAG catalogues [32], and the Old Woman Creek wetland collection [33]. The GTDB and OMDB collections also contributed 168 single-amplified genomes (SAGs) (131 and 37, respectively; Supplementary Table S2). For each record, we retained its genome type and collection-specific provenance and metadata where available (Supplementary Table S2). Duplicate genomes, within or across collections, were retained only once, preferentially in GTDB. MAGs binned from metagenome assemblies that are also part of BGC Atlas were retained deliberately, as their BGCs then provide both sample and genomic context; 287,738 BGCs of MAGs also occur, with identical sequence, in the metagenome assembly from which the MAG was binned.

To apply uniform quality criteria across collections, we assessed all MAGs, SAGs and isolate genomes with CheckM2 [34], using our own CheckM2 v1.1.0 runs for all genomes outside GTDB and the CheckM2 estimates published with GTDB release 232 for GTDB genomes. Following the MIMAG standard [35], genomes with an estimated completeness below 50% or contamination of 10% or more were classified as low quality and excluded together with their BGCs (Supplementary Table S2). CheckM2 completeness and contamination are reported for each genome in the web application.

### 2.2 BGC detection

All metagenome assemblies and genomes (MAGs, SAGs and isolate genomes) were analysed with metaSMASH [36], a scalable implementation of antiSMASH 8.0 [37]. We enabled extended cluster detection and annotation modules with the following parameters:

~~~
–clusterhmmer –asf –cc-mibig –cb-subclusters
–cb-knownclusters –cb-general –pfam2go –rre
–tfbs –tigrfam
~~~

Genes were predicted with Prodigal [38]. Metagenome assemblies were analysed in metagenomic mode (`prodigal-m`), whereas MAGs, SAGs and isolate genomes were analysed in single-genome mode (`prodigal`).

Throughout this work, each antiSMASH region is counted as one BGC; we use the term “region” only where referring to antiSMASH-specific properties, such as region types or contig-edge flags. antiSMASH assigns a product type and a product category (NRPS, PKS, RiPP, terpene, saccharide or other) to each protocluster; a BGC whose protoclusters belong to several categories carries all of them, and summaries by product category therefore count product assignments rather than BGCs. Terpene regions were further divided by product type into canonical terpene and terpene-precursor regions, a region type introduced in antiSMASH 8.

For comparison with characterized BGCs, we considered a BGC similar to a reference in the Minimum Information about a Biosynthetic Gene cluster (MIBiG) database, version 4.0 [39], if KnownClusterBlast, which reports the percentage of genes of a reference cluster that have a significant BLAST hit in the query region, reported a similarity of ≥50%.

After the removal of duplicate and low-quality genomes, BGC Atlas v2 retains 16,109,198 BGCs, of which 12,221,853 derive from metagenome assemblies and 3,887,345 from MAGs, SAGs and isolate genomes.

### 2.3 Gene cluster family inference

We clustered the resulting BGCs into gene cluster families (GCFs) using a seed-and-propagate strategy (Supplementary Figure S6) based on BiG-SLiCE 2.0 [40]. MIBiG 4.0 reference BGCs were included in the BiG-SLiCE clustering to identify families containing a characterized member. We considered a BGC complete if antiSMASH did not flag it as extending to a contig edge. To select the seed set, we additionally derived product-category-specific length baselines from complete, single-protocluster BGCs of GTDB genomes: for each product category, the baseline is the median length of these reference regions (Supplementary Table S3). Because an antiSMASH region comprises the biosynthetic core together with a rule-specific flanking neighbourhood, an incomplete region at least as long as a typical complete region of its category is likely to retain its core and most of its flanking genes. The seed set therefore comprised all complete BGCs together with incomplete BGCs of at least the category-specific median length (3,172,706 BGCs). The length baselines were used only for this selection and do not affect the completeness statistics reported elsewhere. We removed exact duplicates using SHA-256 hashes and clustered the remaining seed regions at 95% sequence identity and 80% bi-directional coverage with MMseqs2 Linclust [41], yielding 1,046,602 representatives. As the representative of each cluster, we selected the member not flagged as extending to a contig edge, if available, and otherwise the longest member, so that each representative is the most complete available sequence of its group. BiG-SLiCE clustered these representatives at a feature-distance threshold of 0.4, yielding 373,816 GCFs, of which 373,360 contained at least one BGC of the collection (the other 456 contained only MIBiG reference BGCs) and formed the seed GCFs of the GCF catalogue.

The remaining 12.94 million non-seed regions were assigned to these GCFs through complementary sequence- and feature-based comparisons. First, MMseqs2 `easy-search` assigned regions by sequence containment when they matched a representative at ≥90% sequence identity and ≥80% query coverage, assigning 54.9% of non-seed regions. Regions not assigned by containment were queried against GCF centroids (protein-domain feature profiles summarizing each family) with BiG-SLiCE and assigned at a nearest-centroid feature distance of ≤0.4, a further 10.1% of non-seed regions.

Because these assignment routes compare regions only with established families, we separately clustered the remaining partial regions using BiG-SLiCE, and assigned the regions left over from this step to the resulting families using the same containment and query-mode procedures. Only BGCs of at least 30% of the category-specific median length founded these families; shorter fragments were only assigned to existing families (Supplementary Table S3). This yielded 426,685 fragmented-region families, of which the 256,512 families with two or more members were added to the GCF catalogue; the 170,173 single-member fragmented-region families were not listed. Seed families were listed regardless of size, including 109,342 single-member families, because a single complete or near-full-length BGC can represent a unique pathway, whereas a single fragment provides too little evidence to define a family. In total, the catalogue contains 629,872 GCFs representing 88.3% of all BGCs (71.9% in seed families and 16.4% in fragmented-region families). The remaining 1,880,410 BGCs consist of 1,710,237 BGCs (10.62%) that were not assigned to any family and 170,173 BGCs (1.06%) in the unlisted single-member fragmented-region families.

### 2.4 Taxonomic assignment

Taxonomic origin was assigned to all BGCs using MMseqs2 `easy-taxonomy` (v18, with the 2bLCA algorithm, sensitivity -s 4 and a maximum query length of 10^6^ bp) [42] against a reference proteome database derived from GTDB release 232 [43]. For BGCs deriving from the GTDB high-quality collection, the taxonomic assignment was taken directly from the source genome. All other BGCs, including those from MAGs, SAGs and isolate genomes of other collections, were assigned individually. Genome-level classifications provided by the other collections were not used, because they are based on different GTDB releases; assigning all remaining BGCs with the same method and reference ensures consistent taxonomy across data sources. For the 1,636,396 BGCs from GTDB genomes that are not part of the reference database, BGC-level assignments agreed with the genome taxonomy for 99.9% of BGCs at phylum, 98.7% at genus and 96.6% at species level whenever an assignment was made; 17.4% and 62.6% of these BGCs remained unassigned at genus and species level, respectively. Overall, 99.4% of BGCs received a taxonomic assignment; 54.2% were resolved to species and 82.1% to genus or a more specific rank.

### 2.5 Harmonization of environmental metadata

All samples, irrespective of source, were mapped to two shared controlled vocabularies: the five-level GOLD ecosystem path [44] and the Environment Ontology (ENVO) biome, environmental feature and environmental material triad [45]. Geographic coordinates, collection dates and host annotations were harmonized alongside these environmental terms.

GOLD paths and other environment classifications provided by the sources (MGnify biomes, the habitat categories of genome catalogues, or a single habitat for collections from one environment) were translated to the GOLD ecosystem classification with fixed mappings, rewriting path segments and standardizing casing, so that records from different sources used the same conventions. For samples without such a classification, we inferred a GOLD path using a priority-ordered, rule-based evidence stack. In order of precedence, this drew on MIxS environmental packages, ENVO biome, feature and material fields, isolation-source and host fields, and, for GTDB genomes, species-level habitat traits from the Madin database [46]. Mapping rules were anchored to specific terms, evaluated from specific to general, and did not relocate records across top-level ecosystem branches; ambiguous records were retained at a more general level rather than assigned to a specific habitat. The rule and source responsible for every inferred value were recorded alongside the annotation, allowing each automated assignment to be audited. After harmonization, all but 28 of the 67,067 metagenomic samples and 94.8% of genomes carried a GOLD ecosystem path, covering 98.4% of BGCs; this includes 94.0% of GTDB genomes, which lack ecosystem annotations in their source. Paths inferred from sample metadata account for 27.4% of metagenomic samples and 20.0% of genomes (39.3% of BGCs); genomes without a GOLD path mostly lack usable sample metadata, and the remainder carry host or isolation-source values not covered by the rules.

### 2.6 Web application

The redesigned web application organizes the resource around interconnected pages for BGCs, GCFs, samples, genome collections and studies (Figure 1). An interactive map shows the geographic distribution of samples, coloured by environment and scaled by BGC count (Figure 1A). BGC pages combine antiSMASH annotations with completeness, GCF membership, taxonomy and source information, and include an interactive gene-cluster view rendered with the BGC-Viewer web components [47]. GCF and study pages summarize biosynthetic composition (Figure 1C) and taxonomic, environmental and geographic distributions. Sample and genome pages connect these records to their assemblies, metadata and originating studies. Identifiers link to relevant external resources, and processed data and sequence archives are available through the download section. A feedback form integrated into the web application allows users to report problems and suggest improvements.

**Figure 1.**
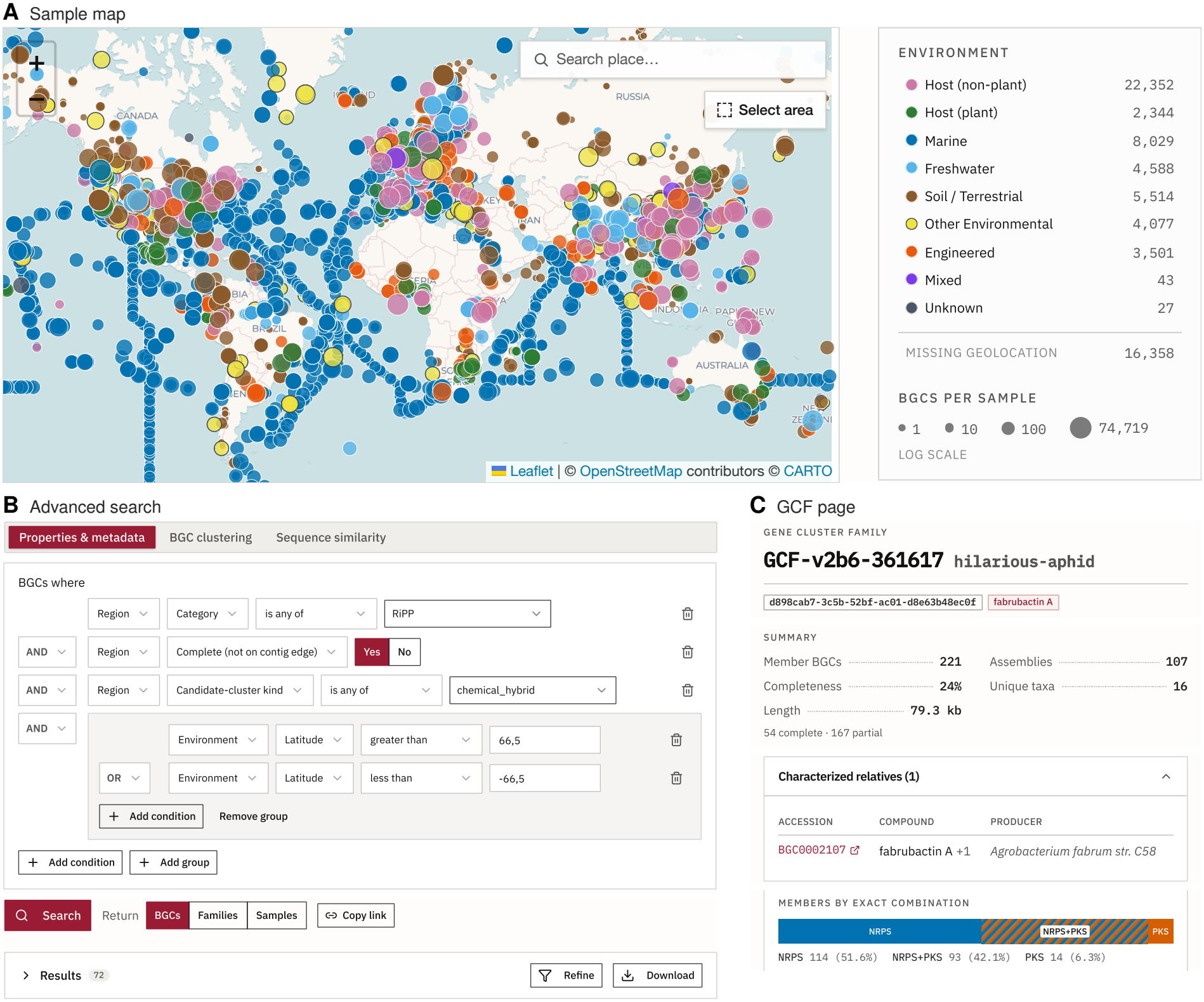
The BGC Atlas v2 web application. (**A**) Interactive map of sample origins, coloured by environment and scaled by the number of BGCs per sample. (**B**) Advanced search with its three modes (properties and metadata, BGC clustering, sequence similarity), showing the query builder with nested Boolean conditions and an example query for complete hybrid RiPP clusters from polar latitudes. (**C**) Page of GCF-v2b6-361617, a family of 221 BGCs, related to fabrubactin A (MIBiG BGC0002107), with summary statistics and members by category combination.

Taxonomic and environmental context is available throughout the application where applicable. Tables can be filtered by taxonomic units, GOLD ecosystems, ENVO terms and geographic location, while accompanying charts and maps update to represent the selected records. This allows users to move directly between an individual BGC, related families and the organisms and environments in which they occur.

### 2.7 Searching and filtering

The application provides four complementary search modes. First, browse tables can be filtered directly by properties including product category, completeness, data source, study, taxonomy, environment and geography. Second, an advanced query builder supports nested Boolean combinations of biosynthetic, functional, taxonomic, environmental and GCF properties (Figure 1B). Available fields include BGC architecture, biosynthetic domains, tailoring enzymes, non-ribosomal peptide synthetase (NRPS) and polyketide synthase (PKS) annotations, features of ribosomally synthesized and post-translationally modified peptides (RiPPs), and similarity to characterized clusters. Search results are accompanied by feature counts, distribution plots and geographic summaries, and can be regrouped by GCF or sample.

Queries are encoded in the URL so that they are persistent and can be shared and revisited. The resulting set of BGCs or GCFs can be exported as a table, and the corresponding sequences and regions can be downloaded in FASTA or GenBank formats (up to 150,000 records per export). Analyses of larger result sets are supported by the complete database tables, indexes and sequence archives provided in the download section.

Third, user-supplied BGCs can be assigned to the gene cluster families of the collection. antiSMASH-compatible GenBank files are uploaded and matched against the GCF catalogue with BiG-SLiCE [40], as in the first release, so that a newly identified cluster can be placed among its closest relatives in the atlas.

Fourth, sequence-based searches allow user-supplied sequences to be queried with DIAMOND [48] against the amino acid sequences of the coding sequences of BGCs in BGC Atlas. Searches run with DIAMOND v2.2.2 (blastp; E-value ≤10^−5^, ≥50% identity, ≥80% query coverage, up to 1,000 hits per query) against all 166.8 million BGC-encoded proteins. A single protein typically completes within 2–4 min and a maximum-size submission of 100 sequences within about 1 h. Search results link back to the corresponding BGC, GCF and sample pages. Result tables can be exported, and larger sequence and database files are provided through the download section.

## 3 Results

### 3.1 Database content

BGC Atlas v2 substantially expands both the scale and the types of sequence data represented in the database. It contains 16,109,198 BGCs identified in 85,340 metagenome assemblies and 750,901 MAGs, SAGs and isolate genomes, representing 44,268 studies across 18 data sources. Studies comprise one or more samples; the 85,340 metagenome assemblies derive from 67,067 metagenomic samples, as a sample can have several assemblies, whereas each MAG, SAG and isolate genome is linked to its own sample record and to the study that reported it. 12.4% of BGCs share their sequence with at least one other BGC; identical BGCs found in different samples, studies, sources or genomes are retained as separate records but are collapsed before GCF clustering. In total, 2,699,040 BGCs (16.8%) are complete, and complete and partial BGCs together are organized into 629,872 gene cluster families. Compared with the first release, which contained 1,854,079 BGCs from 31,316 metagenome assemblies, v2 therefore not only increases the size of the collection more than eightfold, but also adds genome-resolved and isolate data and contains a substantially higher proportion of complete BGCs (16.8% compared with 8.3%; Table 1). Reanalysis of the first-release assemblies with antiSMASH 8 detected 1.27 million BGCs on contigs on which the first release had identified none, 0.80 million of which are terpene-precursor regions, whereas new metagenome assemblies, MAGs, SAGs and isolate genomes contribute 13.2 million BGCs (82.1% of v2; Supplementary Table S6).

**Table 1.** Contents of the first release of BGC Atlas and of v2.

|  | First release | v2 |
|---|---|---|
| Data sources | 1 | 18 |
| Metagenome assemblies | 31,316 | 85,340 |
| MAGs | 0 | 504,037 |
| Isolate genomes | 0 | 246,696 |
| SAGs | 0 | 168 |
| BGCs | 1,854,079 | 16,109,198 |
| Complete BGCs<sup>2</sup> | 153,278 (8.3%) | 2,699,040 (16.8%) |
| Gene cluster families<sup>1</sup> | 18,566 | 629,872 |
| BGCs with taxonomic assignment | none | 99.4% |

First-release counts are those reported in the original publication [8]; the first release drew all metagenome assemblies from MGnify.
<sup>1</sup> Family counts are not directly comparable between releases: the first release clustered with BiG-SLiCE 2.0 at a fixed threshold and assigned most BGCs by search against family models, whereas v2 clusters de novo with the seed-and-propagate procedure described in Methods.
<sup>2</sup> Completeness is not strictly comparable between releases either: v2 was analysed with antiSMASH 8, which introduced terpene-precursor regions. Excluding regions annotated only as terpene-precursor, 16.1% of v2 BGCs (2,012,679 of 12,539,567) are complete.

RiPPs form the largest category among the 16.5 million product assignments of the v2 collection (28.5%), followed by terpene-precursor (21.9%), terpene (14.7%), NRPS (13.9%) and PKS (11.5%). Two factors contribute to the differences in biosynthetic composition between v1 and v2. First, terpene-precursor regions are a new region type introduced in antiSMASH 8 and are reported separately from canonical terpene BGCs. When the v1 assemblies were re-analysed, 796,307 terpene-precursor regions were detected on contigs in which v1 had identified no BGC. Excluding these regions, the composition of v2 is considerably more similar to that of v1. Second, v2 draws far more evenly on environmental and host-associated samples than v1 did (48.0% and 42.7% of BGCs, against 22.3% and 66.8%), shifting the collection away from the RiPP-rich host-associated composition that dominated v1. Despite the increased proportion of complete BGCs, assembly fragmentation remains substantial, with 83.2% of detected BGCs extending to a contig edge.

This fragmentation is also reflected in the GCF catalogue, which contains 373,360 seed families and 256,512 fragmented-region families. Overall, 14,228,788 BGCs (88.3%) are assigned to a GCF. Because BiG-SCAPE 2.0 [40] could not be run on collections of this size, and BiG-SLiCE 2.0 [40] would have been impractical on the full collection (Supplementary Table S5); we developed a seed-and-propagate GCF inference. It nevertheless performed on par with both tools on curated datasets of complete BGCs and, on fragmented BGCs, agreed more closely with BiG-SCAPE than BiG-SLiCE did (Supplementary Section S1). Only 1,328 GCFs clustered with an MIBiG 4.0 reference BGC during BiG-SLiCE clustering, while 944,335 BGCs (5.9%) showed at least 50% similarity to an MIBiG reference in KnownClusterBlast, spanning 62,114 GCFs (9.9% of all GCFs), reflecting the more permissive, gene-level nature of this measure. Thus, even within this very large collection, only a relatively small fraction of GCFs is closely connected to experimentally characterized biosynthetic pathways.

A major addition in v2 is the ability to place this biosynthetic diversity into taxonomic as well as environmental context. Taxonomic assignments are available for 99.4% of BGCs, with 82.1% resolved to genus and 54.2% to species level. In total, the collection spans 197 phyla, 31,800 genera and 160,634 species. GOLD ecosystem paths are available for 95.2% of samples, covering 98.4% of BGCs, and ENVO biome terms for 94.4% of samples, covering 97.4%. Geographic coordinates are available for 75.6% of metagenome samples, 91.6% of MAG, 64.9% of SAG records, and 25.0% of isolate records. Together, these annotations allow individual BGCs and GCFs to be examined in relation to both the organisms that encode them and the environments in which they occur.

### 3.2 Biosynthetic composition differs between ecosystems

The environmental context available in BGC Atlas v2 makes it possible to ask how the biosynthetic composition represented in the database differs between ecosystems. Clear differences were already apparent at the highest level of the GOLD hierarchy (Figure 2A). Terpenes accounted for 23.6% of product assignments in environmental but only 4.9% in host-associated samples, whereas RiPPs showed the opposite pattern, accounting for 19.0% and 39.6%, respectively. Engineered samples were intermediate, and terpene-precursor regions occurred at broadly similar frequencies across ecosystem types. These differences continued at finer environmental scales. At the next level of the GOLD hierarchy, aquatic samples contained the highest proportion of terpene BGCs and human-associated samples the highest proportion of RiPPs, while NRPSs were most abundant in plant-associated, built-environment and terrestrial samples (Supplementary Figure S1).

**Figure 2.**
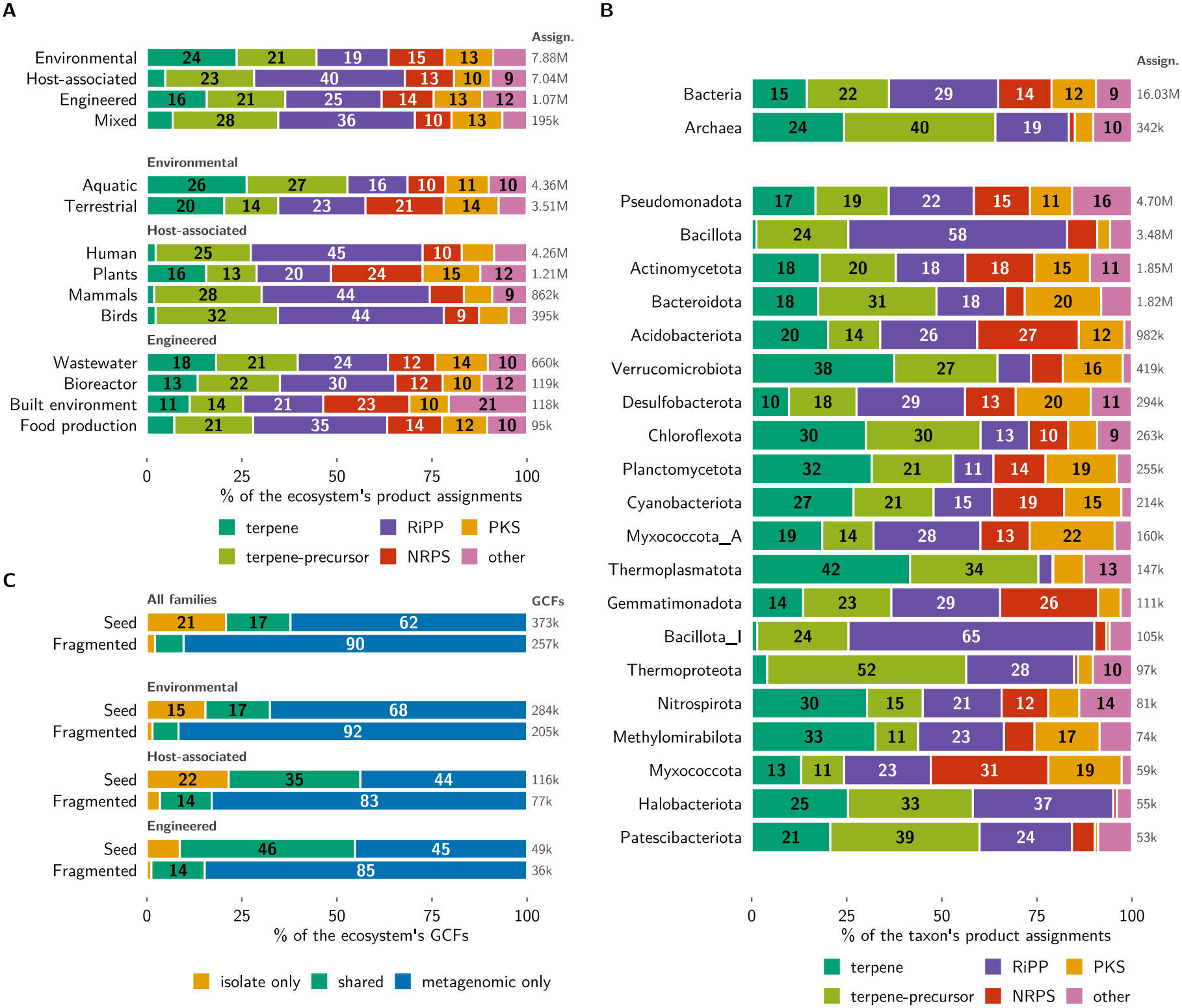
Biosynthetic composition across environments and taxa, and sharing of gene cluster families between sequence sources. In (**A**) and (**B**), bars show the proportions of product assignments within each group, with total numbers of assignments at right. Following antiSMASH, a BGC whose protoclusters belong to more than one product category (2.1% of BGCs) contributes one assignment to each of these categories. (**A**) The four top-level GOLD ecosystems and the ten largest level-2 ecosystems, grouped by parent ecosystem. Ecosystem counts accumulate up the hierarchy and include samples annotated only at higher levels. (**B**) The two domains and the 20 largest GTDB phyla. Rows are ordered by number of assignments within each block. Canonical terpene and terpene-precursor regions are shown separately. (**C**) Proportions of isolate-only, shared and metagenomic-only GCFs overall and within the Environmental, Host-associated and Engineered ecosystems, shown separately for seed GCFs and fragmented-region GCFs (see Methods). Further stratifications are shown in Supplementary Figure S4. Metagenomic sources comprise metagenome assemblies, MAGs and SAGs.

Similar contrasts were apparent among individual habitats (Supplementary Figure S2): aquatic habitats were consistently terpene-rich, whereas soil and plant-associated habitats contained higher proportions of NRPSs. Biosynthetic composition also differed among habitats belonging to the same host. Nearly half of BGCs from the human digestive system were RiPPs, whereas skin, respiratory and circulatory samples contained roughly threefold more NRPSs than the digestive system.

Because metagenomic assemblies differ substantially in fragmentation, we repeated the analysis using complete clusters only. The strong contrast between environmental and host-associated samples persisted, including terpene enrichment in soil and marine habitats and RiPP enrichment in the human digestive system. Differences among environmental habitats were reduced, suggesting that fragmentation affects some finer-scale comparisons but does not explain the principal ecosystem-level patterns.

### 3.3 Biosynthetic composition varies across and within lineages

The taxonomic assignments added in v2 also make it possible to examine how biosynthetic composition varies across microbial lineages (Figure 2B). Archaea were enriched in terpene and terpene-precursor regions and depleted in NRPS and PKS BGCs relative to Bacteria; most notably, NRPSs represented only 1.6% of archaeal product assignments compared with 14.0% in Bacteria.

Differences among major phyla were even more pronounced. Thermoplasmatota and Verrucomicrobiota were terpene-rich, Bacillota and Bacillota_I were dominated by RiPPs, Myxococcota, Acidobacteriota and Gemmatimonadota were enriched in NRPSs, and Myxococcota_A, Bacteroidota and Desulfobacterota showed the highest proportions of PKSs. NRPS depletion was consistent across the major archaeal phyla, indicating a broader domain-level pattern rather than an effect of a single lineage.

These lineage-specific patterns also help explain some of the environmental differences described above. Bacillota dominated host-associated BGCs, whereas environmental BGCs were largely contributed by Pseudomonadota, Actinomycetota and Acidobacteriota. However, habitat-associated differences remained within individual phyla. Environmental Pseudomonadota and Bacteroidota were more terpene-rich than their host-associated counterparts, host-associated Bacillota contained more RiPPs, and soil Pseudomonadota contained more than twice the proportion of NRPS BGCs found in marine samples. The differences between habitats therefore reflect both changes in which taxa are represented and differences in biosynthetic composition within the same broad lineages.

### 3.4 Gene cluster families are unevenly shared between isolate genomes and metagenomic data

By combining isolate genomes with metagenomic assemblies, MAGs and SAGs, BGC Atlas v2 also makes it possible to ask how much of the biosynthetic diversity recovered from culture-independent sequence data is represented among isolate genomes. Most GCF diversity was not. Among seed GCFs, which are anchored by complete or near-full-length BGCs, 62.2% occurred only in metagenomic assemblies, MAGs or SAGs, whereas 17.0% were shared between these sources and isolate genomes (Figure 2C). This proportion was similar when only complete BGCs were considered (61.2%; Supplementary Figure S4). Fragmented-region GCFs were almost exclusively metagenomic (90.3%), as expected because fragmented BGCs derive mainly from metagenomic data, and their number should be regarded as an upper bound on the number of distinct families, since fragments of one pathway may fall into different families. Shared families nevertheless contained 62.4% of the BGCs assigned to a GCF (Supplementary Figure S3A; 71.9% of those in seed GCFs), indicating that the overlap between isolate and metagenomic sources is concentrated among larger, more frequently recovered GCFs, whereas much of the remaining family-level diversity is source-specific.

The extent of this pattern differed between ecosystems: metagenomic-only families made up 67.6% of seed GCFs in environmental samples, but fewer than half in host-associated (43.8%) and engineered (45.2%) samples, in which a larger share of families was also found in isolate genomes (34.6% and 46.1%, respectively). The extent of overlap also varied among biosynthetic classes: isolate-only families were relatively uncommon among terpene and RiPP GCFs but more frequent among NRPS and PKS GCFs (Supplementary Figure S3C).

## 4 Discussion

BGC Atlas v2 expands both the underlying data and the ways in which bacterial and archaeal biosynthetic diversity can be explored. Broader sampling of metagenomes reduces the strong overrepresentation of host-associated samples in the first release: environmental and host-associated samples now contribute similar numbers of BGCs. Together with the terpene-precursor regions introduced in antiSMASH 8, this change in sampling explains much of the lower proportion of RiPP BGCs in v2 relative to v1. At the same time, the redesigned web application introduces new search modes and filters across studies, samples, genomes, BGCs and GCFs. BGC Atlas v2 therefore represents an expansion not only in scale but also in scope, allowing BGCs to be linked to organisms and explored jointly by taxonomy, environment and biosynthetic features.

Despite this expanded coverage, BGC Atlas v2 also reflects important limitations of the public sequence data and metadata on which it is built. The environmental component of the database remains dominated by short-read metagenomes, for which fragmented assemblies can truncate BGCs and complicate their detection, GCF assignment and taxonomic classification. This effect is particularly clear in the Microflora Danica project, for which both short- and long-read MAG catalogues are available [20, 32]. In our analysis of these catalogues, 81.5% of the 94,322 BGCs detected in 15,410 long-read MAGs were complete, compared with only 2.3% of the 131,720 BGCs in 19,161 short-read MAGs (Supplementary Table S2). This difference persisted when the comparison was restricted to the 99 samples represented in both catalogues (81.7% and 4.0%, respectively). Although the inclusion of long-read metagenomes, MAGs and isolate genomes improves the representation of complete clusters, these data remain unevenly distributed across taxa and environments.

The available metadata are similarly heterogeneous in completeness, accuracy, detail and formatting. Harmonization improves comparability but cannot recover information that was never reported, and inferred annotations remain limited by the quality of the original metadata. Taxonomic assignments inferred from individual BGCs are also affected by horizontal gene transfer. Because BGCs are frequently transferred between lineages, and short or chimeric contigs carry limited or conflicting phylogenetic signal, these assignments reflect the closest reference lineages of a BGC and may not always identify the organism that carries it. The lowest-common-ancestor approach mitigates this by assigning BGCs with conflicting signal to higher taxonomic ranks, and assignments at low ranks should be interpreted with this caveat in mind.

The expanded environmental and taxonomic context nevertheless reveals clear structure in the biosynthetic diversity represented in the collection. Biosynthetic composition differs both between habitats and between microbial lineages, and these two axes are only partly aligned. Much of the contrast between environmental and host-associated samples along a terpene–RiPP axis follows from differences in taxonomic composition, particularly the dominance of Bacillota in host-associated samples. However, the same environmental contrasts are also observed within Pseudomonadota, Bacteroidota and Bacillota, while NRPS clusters are concentrated both in particular habitats and in lineages such as Myxococcota, Acidobacteriota and Gemmatimonadota. Thus, the biosynthetic potential represented in microbial communities reflects both which taxa are present and how biosynthetic repertoires vary within those taxa. These patterns describe the current BGC Atlas collection and should not be interpreted as unbiased estimates of BGC prevalence across all natural environments, given the uneven sampling of available sequence data.

The inclusion of both isolate genomes and culture-independent sequence data also highlights how much biosynthetic diversity is missed when analyses are restricted to cultivated organisms. Even when only seed GCFs are considered, more than 60% (62.2%) were recovered only from metagenomic assemblies, MAGs or SAGs. The GCFs shared between isolate and metagenomic sources contained many of the most frequently recovered BGCs, whereas much of the remaining family-level diversity was restricted to culture-independent data.

The continued growth in the number of detected BGCs also makes GCF inference an increasingly important computational challenge. GCFs provide a practical way to organize related BGCs and to compare the novelty and distribution of biosynthetic pathways across ecosystems and taxa. The seed-and-propagate strategy developed here enabled the current collection to be clustered at scale by combining seed families with complementary sequence- and feature-based assignments. However, as BGC collections continue to expand in both size and heterogeneity, maintaining biologically meaningful groupings will become progressively more demanding. GCF inference is therefore likely to remain a major bottleneck for future updates of BGC Atlas and other large-scale BGC resources.

At the same time, this continued growth is revealing an extraordinary breadth of prokaryotic biosynthetic potential. Even a collection of more than 16 million predicted BGCs resolves into hundreds of thousands of GCFs, only a small fraction of which are closely connected to experimentally characterized pathways. BGC Atlas v2 makes this diversity accessible through a web application that allows individual BGCs to be placed into a global taxonomic and environmental context and large-scale patterns to be explored across the collection. The scale and diversity of the current resource also make clear that we are still far from capturing the full biosynthetic potential encoded in bacterial and archaeal genomes and microbial communities. We expect BGC Atlas to continue to support both the prioritization of biosynthetic novelty for natural-product discovery and broader investigations into the ecology and evolution of secondary metabolism.

## Abbreviations

BGC: biosynthetic gene cluster
ENVO: Environment Ontology
GCF: gene cluster family
GOLD: Genomes OnLine Database
GTDB: Genome Taxonomy Database
MAG: metagenome-assembled genome
MIBiG: Minimum Information about a Biosynthetic Gene cluster
NRPS: non-ribosomal peptide synthetase
OMDB: Ocean Microbiomics Database
PKS: polyketide synthase
RiPP: ribosomally synthesized and post-translationally modified peptide
SAG: single-amplified genome

## 5 Supplementary data

Supplementary data are available alongside this preprint.

## 6 Acknowledgements

The authors acknowledge support from the High Performance and Cloud Computing Group at the Zentrum für Datenverarbeitung of the University of Tübingen, the state of Baden-Württemberg through bwHPC, and the German Research Foundation (DFG) under project number 455787709 (bwForCluster BinAC 2). We thank the Cluster of Excellence Controlling Microbes to Fight Infection (CMFI), funded by the DFG under Germany’s Excellence Strategy – EXC 2124 – 390838134, for structural support. We also thank Luděk Sehnal for extensively testing the resource and reporting bugs. This work was supported by the de.NBI Cloud within the German Network for Bioinformatics Infrastructure (de.NBI) and ELIXIR-DE (Forschungszentrum Jülich and W-de.NBI-001, W-de.NBI-004, W-de.NBI-008, W-de.NBI-010, W-de.NBI-013, W-de.NBI-014, W-de.NBI-016, W-de.NBI-022).

## 7 Conflict of interest statement

The authors declare no competing interests.

## 8 Funding

This work was supported by the German Center for Infection Research (DZIF) [TTU Novel Antibiotics 09.726] and the Momentum Program of the Volkswagen Foundation (Volkswagen Stiftung) [0072511-00].

## 9 Data availability

BGC Atlas v2 is freely available at https://bgc-atlas.ziemertlab.com. The download section provides all BGCs as GenBank files and as nucleotide and protein FASTA files, the GCF catalogue and BGC-to-GCF memberships as tab-separated tables, the complete set of database tables in TSV and Parquet formats, a PostgreSQL dump of the core relational tables, and provenance tables linking BGC identifiers to their source assemblies, together with SHA-256 checksums for every file. Data produced by BGC Atlas, comprising BGC annotations, family assignments, taxonomic assignments and harmonized metadata, are released under a CC BY 4.0 license; the underlying assemblies and genomes retain the licenses of their source repositories. All statistics refer to data release R2.0.

The source code of the web application, the data ingestion and metadata harmonization pipeline, the GCF inference pipeline, including the benchmarking scripts, and the scripts for taxonomic assignment and genome quality filtering is available at https://github.com/ZiemertLab/bgc-atlas-web-v2 under the GNU Affero General Public License v3.0 (AGPL-3.0). The version described here is archived at Figshare (https://doi.org/10.6084/m9.figshare.34748379) [49]. BGC detection used metaSMASH [36].

## 10 Author contributions statement

C.B.: Conceptualization, Data curation, Formal analysis, Investigation, Methodology, Project adminis-tration, Software, Validation, Visualization, Writing–original draft, Writing–review and editing. A.T.T.: Investigation, Methodology, Writing–review and editing. N.Z.: Conceptualization, Funding acquisition, Project administration, Supervision, Writing–original draft, Writing–review and editing.

## Notes

### Competing Interest Statement

The authors have declared no competing interest.

### Summary of Updates

BGC Atlas v2 is now hosted at https://bgc-atlas.ziemertlab.com. The source code version described in the paper is archived at Figshare (doi:10.6084/m9.figshare.34748379), and the data release is referred to as R2.0. Other changes are minor: a citation to Supplementary Figure S3A was added, and the Acknowledgements were extended (CMFI, de.NBI Cloud/ELIXIR-DE, and Luděk Sehnal).

