## Supplementary Material for "BGC Atlas v2: biosynthetic gene clusters with taxonomic and environmental context at scale"

**Supplementary Table S1:** Metagenome assemblies in BGC Atlas v2, by source. Complete BGCs are those not flagged as extending to a contig edge by antiSMASH, as in Table 1 of the main text. Metalog, which supplied sample metadata for SPIRE assemblies, is not listed as it contributed no sequence data. Genome collections are listed in Supplementary Table S2.

| Source | Assemblies | BGCs | Complete BGCs (%) |
|---|---|---|---|
| MGnify | 66,922 | 6,618,170 | 673,744 (10.2) |
| SPIRE | 18,080 | 5,102,636 | 166,403 (3.3) |
| Microflora Danica | 154 | 370,910 | 188,185 (50.7) |
| Deep-sea hydrothermal deposits | 70 | 35,554 | 2,763 (7.8) |
| FRAM Strait time series | 47 | 1,321 | 602 (45.6) |
| Surface-ocean diel survey | 4 | 20,222 | 2,082 (10.3) |
| Schönbuch forest soil | 2 | 15,413 | 7,053 (45.8) |
| Great Barrier Reef seawater (long-read) | 27 | 38,505 | 16,542 (43.0) |
| Helgoland spring bloom 2020 (long-read) | 34 | 19,122 | 9,430 (49.3) |
| Total | 85,340 | 12,221,853 | 1,066,804 (8.7) |

**Supplementary Table S2:** Genome collections in BGC Atlas v2, by source and genome type. Genomes contained in more than one collection are counted once, and low-quality genomes are excluded (Methods). Complete BGCs are defined as in Supplementary Table S1. Across Supplementary Tables S1 and S2, BGC Atlas v2 comprises 85,340 metagenome assemblies and 750,901 genomes containing 16,109,198 BGCs, of which 2,699,040 (16.8%) are complete.

| Source | Genomes | Isolates | MAGs | SAGs | BGCs | Complete BGCs (%) | Low quality[a] |
|---|---|---|---|---|---|---|---|
| GTDB high-quality genomes | 250,745 | 233,506 | 17,108 | 131 | 1,918,472 | 1,264,778 (65.9) | 25 |
| OMDB | 242,119 | 2,357 | 239,725 | 37 | 773,016 | 114,019 (14.7) | 26,212 |
| SPIRE | 102,496 | 10,833 | 91,663 | 0 | 412,524 | 78,041 (18.9) | 418 |
| GEM | 51,506 | 0 | 51,506 | 0 | 184,780 | 56,817 (30.7) | 977 |
| SMAG | 36,258 | 0 | 36,258 | 0 | 224,873 | 23,822 (10.6) | 2,993 |
| TPMC | 30,960 | 0 | 30,960 | 0 | 138,089 | 14,673 (10.6) | 1,229 |
| Microflora Danica (short-read) | 19,161 | 0 | 19,161 | 0 | 131,720 | 3,003 (2.3) | 92 |
| Microflora Danica (long-read) | 15,410 | 0 | 15,410 | 0 | 94,322 | 76,850 (81.5) | 179 |
| Old Woman Creek wetland | 2,246 | 0 | 2,246 | 0 | 9,549 | 233 (2.4) | 262 |
| Total | 750,901 | 246,696 | 504,037 | 168 | 3,887,345 | 1,632,236 (42.0) | 32,387 |

[a] Genomes excluded as low quality according to MIMAG (completeness $<$50% or contamination $\geq$10%; Methods); they are not part of BGC Atlas v2 and not included in the other columns. Two GTDB MAGs without CheckM2 estimates were retained.

**Supplementary Table S3:** Product-category length baselines used for seed selection in GCF inference. Baselines were computed from the 1,078,941 BGCs of GTDB genomes that were not flagged as extending to a contig edge and contained a single protocluster, grouped by the category of their (first) protocluster. Length is the length of the antiSMASH region. Seed BGCs were all BGCs not on a contig edge, together with BGCs on a contig edge of at least the median length (p50). In the fragmented-region layer, only BGCs of at least 0.3×p50 founded new families; shorter fragments were only assigned to existing families. p10 and 0.3×p50 additionally define the length-based completeness classes used in the benchmark (Supplementary Section S1).

| Product category | Reference BGCs | p10 (bp) | p50 (bp) | 0.3×p50 (bp) |
|---|---:|---:|---:|---:|
| RiPP | 323,196 | 10,831 | 21,131 | 6,339 |
| Terpene | 313,570 | 20,867 | 20,900 | 6,270 |
| Other | 237,963 | 11,212 | 22,090 | 6,627 |
| PKS | 109,544 | 41,086 | 41,263 | 12,379 |
| NRPS | 94,370 | 33,972 | 44,320 | 13,296 |
| Saccharide | 298 | 26,502 | 39,743 | 11,923 |

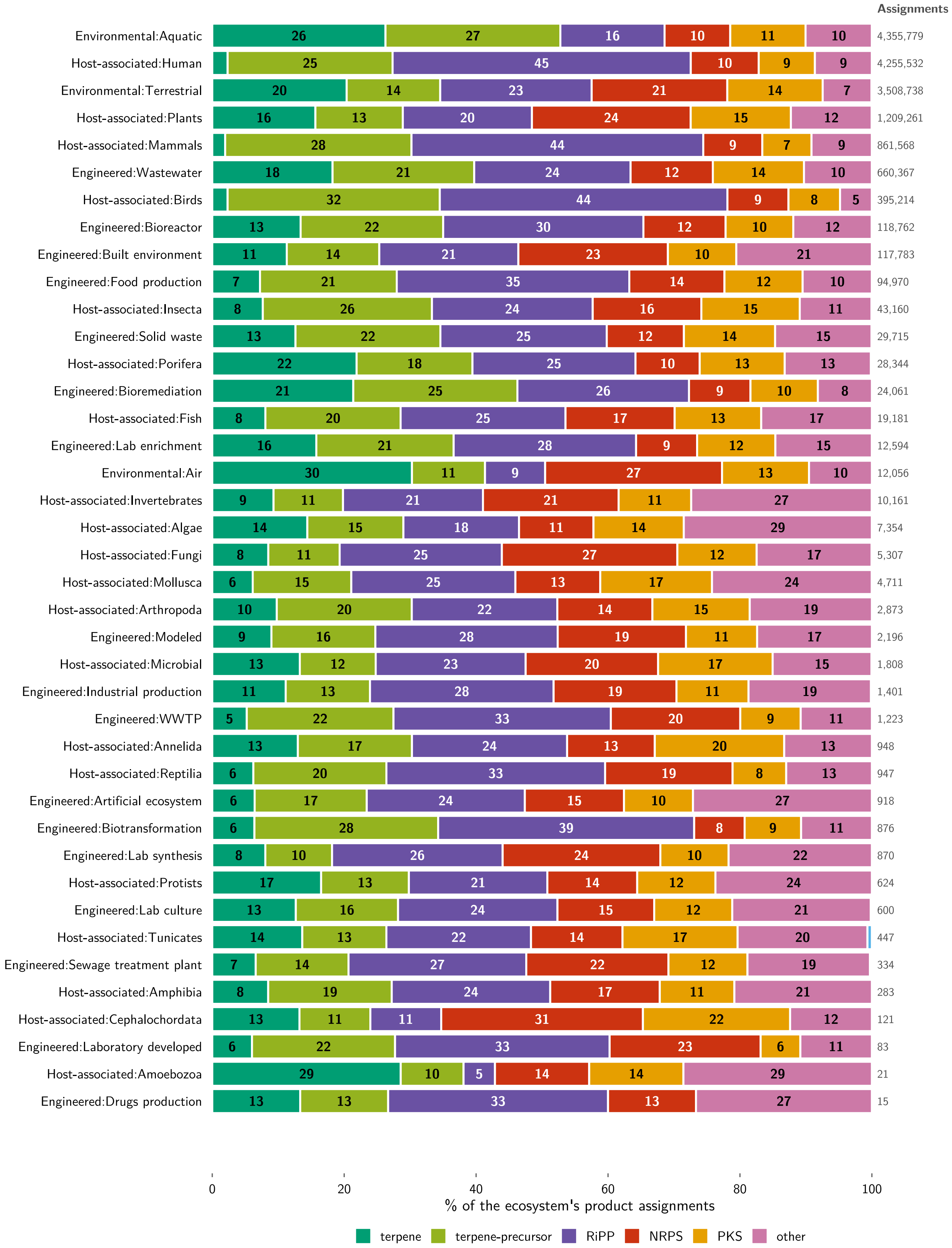


**Supplementary Figure S1:** Biosynthetic composition of BGCs across all 42 level-2 GOLD ecosystems. Each bar shows the distribution of predicted product categories within one ecosystem as a percentage of that ecosystem's product assignments, with total numbers of assignments given at the right, and rows are ordered by number of assignments. A BGC whose protoclusters belong to more than one product category contributes one assignment to each of these categories. Small ecosystems at the foot of the panel are based on few samples and their composition should be interpreted accordingly.

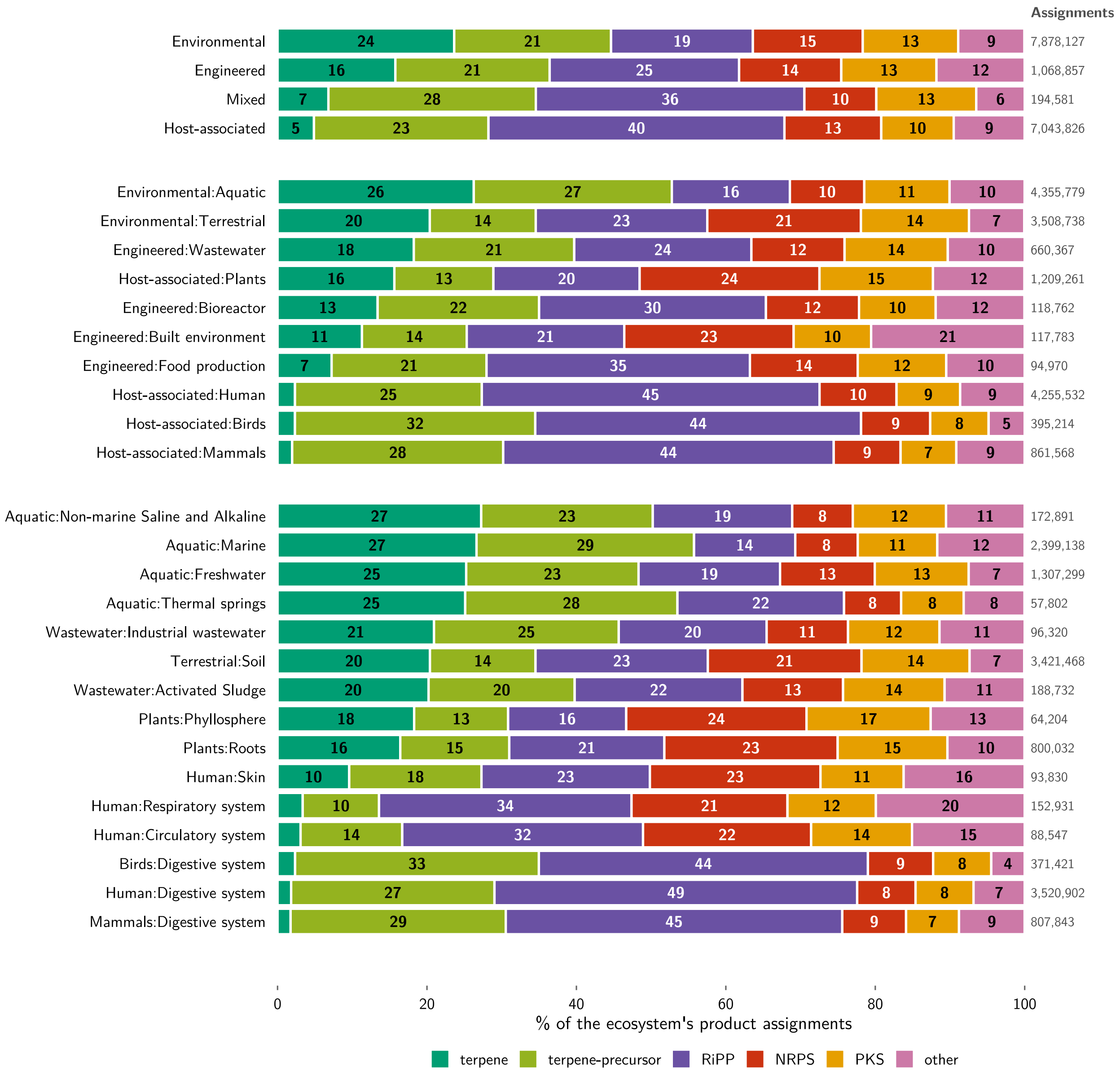


**Supplementary Figure S2:** Biosynthetic composition of BGCs across the GOLD ecosystem hierarchy. Bars, counts and category definitions are as in Supplementary Figure S1. Blocks from top to bottom: the four top-level GOLD ecosystems (Environmental, Host-associated, Engineered and Mixed); the ten largest level-2 ecosystems; and the fifteen largest level-3 ecosystems. Row labels give each ecosystem together with its parent level (level 1:level 2 and level 2:level 3, respectively).

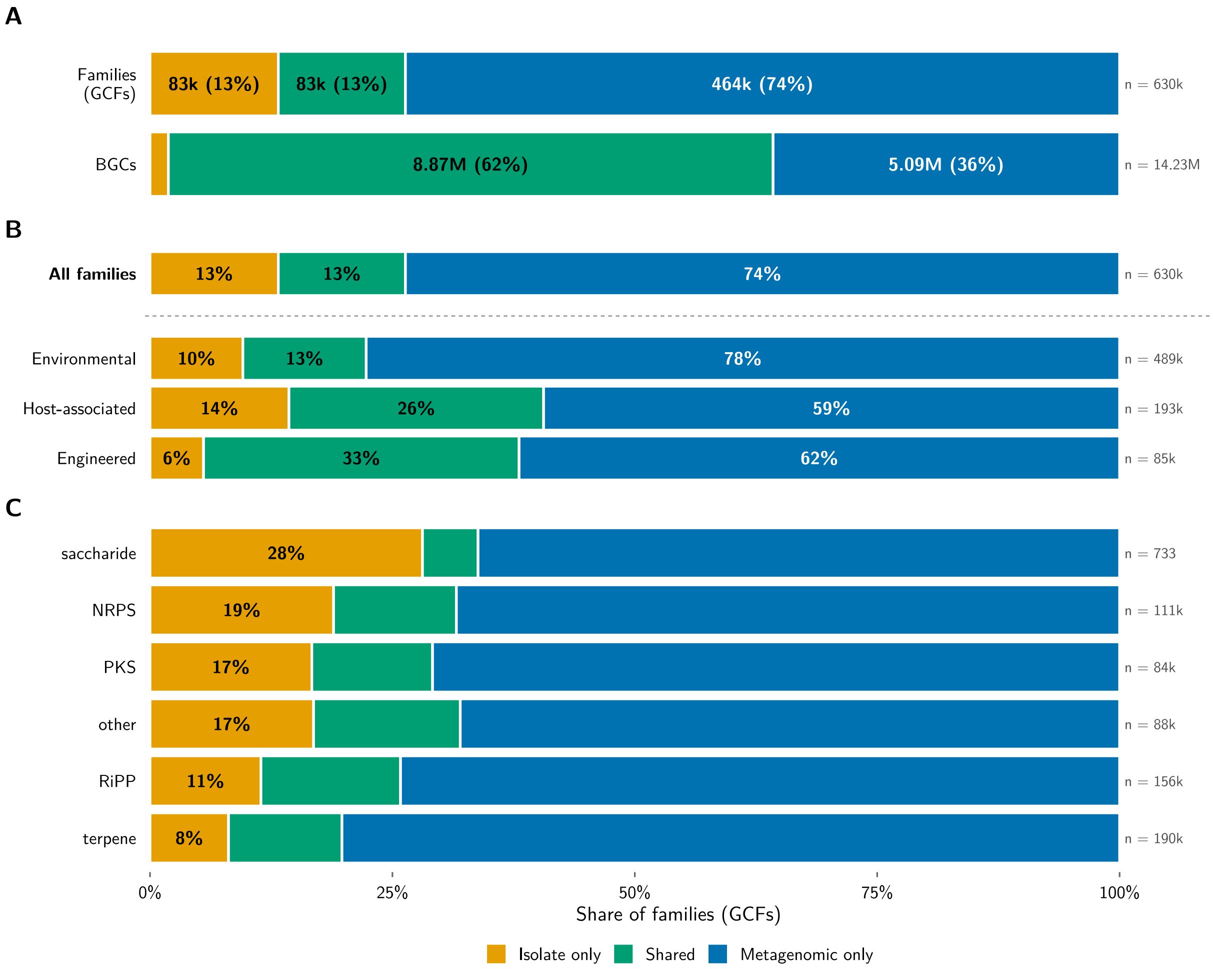


**Supplementary Figure S3:** Sharing of GCFs between isolate genomes and metagenomic sequence sources, in full. (**A**) Family- and BGC-weighted proportions of isolate-only, shared and metagenomic-only GCFs. (**B**) Family-weighted proportions among GCFs represented in the three top-level GOLD ecosystem groups. Ecosystem groups overlap because a GCF may occur in more than one group. (**C**) Family-weighted proportions by dominant biosynthetic product category. Metagenomic sources comprise metagenome assemblies, MAGs and SAGs. Only primary GCF assignments were considered. Panels (**A**)–(**C**) include both seed and fragmented-region GCFs; Figure 2C of the main text and Supplementary Figure S4 show them separately.

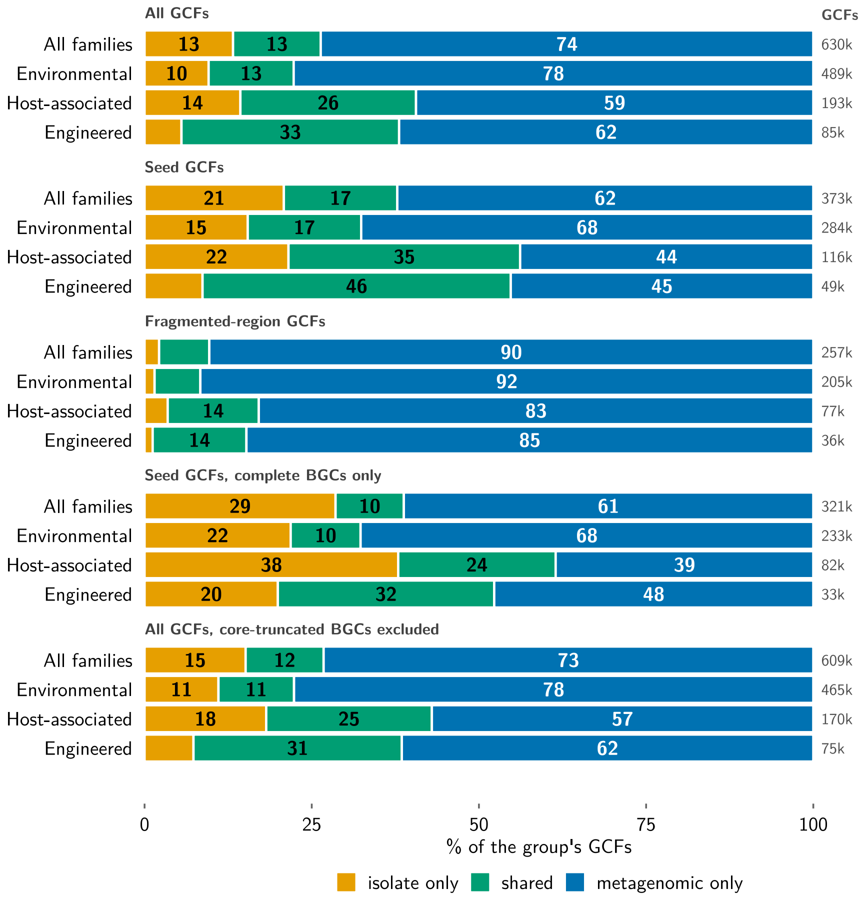


**Supplementary Figure S4:** Sharing of GCFs between isolate genomes and metagenomic sequence sources, stratified by family type and BGC completeness. Each block shows family-weighted proportions of isolate-only, shared and metagenomic-only GCFs overall and within the three top-level GOLD ecosystem groups (rows overlap because a GCF may occur in more than one group). From top to bottom: all GCFs, as in Supplementary Figure S3B; seed GCFs only; fragmented-region GCFs only; seed GCFs, with sharing and ecosystems determined from complete BGCs (not flagged as extending to a contig edge) only; and all GCFs after removing BGCs whose biosynthetic core reaches a contig end (protocluster core within 10 bp of the contig start or end). Metagenomic sources comprise metagenome assemblies, MAGs and SAGs. Only primary GCF assignments of non-redundant BGCs in listed GCFs were considered.

### Supplementary Section S1: Benchmarking of GCF inference

**Methods compared.** This section evaluates the GCF inference procedure described in the Methods of the main text (seed selection, SHA-256 deduplication, MMseqs2 Linclust at 95% identity, selection of representatives, BiG-SLiCE clustering of representatives together with MIBiG 4.0 at a feature distance of 0.4, sequence containment and BiG-SLiCE query mode for non-seed BGCs, and the fragmented-region layer; Supplementary Figure S6), referred to as "this work" in the tables and figures. It was run unchanged on each benchmark input, except that the product-category length baselines were fixed to the values derived for BGC Atlas v2 (Supplementary Table S3) rather than recomputed. It was compared with BiG-SLiCE 2.0.2 and BiG-SCAPE 2.0.3, each at its default threshold (0.4 and 0.3, respectively). BiG-SLiCE was run on all BGCs of each input. For the curated benchmark, BiG-SCAPE was run with the settings used for the curated benchmark of the original publication (`--mix --classify none --include-singletons --alignment-mode glocal --extend-strategy legacy`); for the collection subset, with the settings used there for antiSMASH-DB (`--record-type protocluster --classify class`, singletons excluded). All runs were performed on a high-performance computing cluster with 96 threads per job and nodes with up to 2 TB of memory.

**Curated benchmark.** We used the curated datasets with expert-assigned GCFs published with BiG-SCAPE 2.0 and BiG-SLiCE 2.0, excluding the fungal *Fusarium* dataset because BGC Atlas v2 covers bacterial and archaeal BGCs only. Scoring followed the benchmark module of BiG-SCAPE 2.0: BGCs assigned to two curated families kept the last assignment, only BGCs labelled by both the curation and the method were scored, and homogeneity, completeness and V-measure were computed with scikit-learn. For datasets annotated with antiSMASH 4, which lack region features, the contig-edge flag was taken from the cluster feature.

**Collection subsets.** No curated ground truth exists for fragmented metagenomic BGCs. We therefore drew three nested subsets of the collection (101,589 $\subset$ 507,518 $\subset$ 1,013,058 BGCs), stratified by redundancy (groups of near-identical BGCs were sampled together), completeness, product category and data source; all marginal distributions were within 1.2 percentage points of those of the full collection. The subsets were drawn before duplicate assemblies and low-quality genomes were removed from the collection (Methods), which together affected about 3% of BGCs. Each method was run on each subset in isolation, and our assignments were additionally compared with those that the same BGCs received in the full collection. Agreement was measured with the adjusted Rand index (ARI), V-measure, homogeneity, completeness and pairwise precision and recall, overall and stratified by length-based completeness class: complete (not on a contig edge and at least the category median length, p50), near-complete (on a contig edge and $\geq$p50), core-complete ($\geq$p10), partial ($\geq$0.3$\times$p50) and fragment ($<$0.3$\times$p50). BiG-SCAPE was run on the 0.1-million subset only; because it assigns families to protoclusters, each BGC was given the family of its first protocluster.

**Scalability.** We extracted the regions of antiSMASH-DB 5 (antiSMASH 8) from its 56,846 genomes, excluding 422 fungal genomes, which yielded 528,998 prokaryotic BGCs.

**Results.** On the curated datasets, our procedure performed on par with both tools at their default thresholds (mean V-measure 0.856, 0.857 and 0.850 for this work, BiG-SLiCE and BiG-SCAPE, respectively; Supplementary Table S4). Because these datasets consist of mostly complete BGCs, our procedure reduces to deduplication followed by BiG-SLiCE clustering here, and the benchmark shows that this preserves BiG-SLiCE's performance. On the collection subsets, agreement with BiG-SLiCE was high for complete BGCs (ARI 0.92–0.96) and decreased with fragmentation (ARI

0.23–0.26 for fragments), consistently across subset sizes (Supplementary Figure S5A). For fragments, our procedure split BiG-SLiCE families (completeness 0.76–0.81) but rarely merged them (homogeneity 0.89), producing more, smaller families. With BiG-SCAPE as an independent reference on the 0.1-million subset (Supplementary Figure S5B), our assignments agreed better with BiG-SCAPE than those of BiG-SLiCE overall (ARI 0.35 and 0.18) and for fragments (ARI 0.30 and 0.09). Of the BGC pairs grouped by BiG-SCAPE, 83.2% were also grouped by our procedure and 61.9% by BiG-SLiCE; of the pairs grouped by each method, BiG-SCAPE also grouped 22.5% for our procedure and 10.2% for BiG-SLiCE (fragments: 18.7% and 5.1%), BiG-SCAPE being the most finely resolved of the three methods (29,819 families, compared with 27,871 and 21,246). Assignments obtained on the subsets in isolation agreed with those in the full collection (V-measure 0.96–0.97; ARI 0.67–0.74), the full collection merging some subset families through additional bridging BGCs; this comparison is restricted to the 97% of subset BGCs that remain in the current release. On antiSMASH-DB 5, our procedure clustered 99.2% of BGCs and agreed closely with BiG-SLiCE (ARI 0.83, V-measure 0.97). Runtimes are given in Supplementary Table S5. BiG-SCAPE could not complete the antiSMASH-DB 5 run: with its default multiprocessing mode, it exceeded the 1.9 TB memory limit after 10.7 h, while comparing an estimated 12.7 billion BGC pairs; at the throughput of its memory-conserving mode, the run would require an estimated 18–24 days, beyond the 14-day limit of the cluster.

**Limitations.** Agreement with BiG-SCAPE on fragmented BGCs is not a measure of accuracy, because no ground truth exists for such BGCs, and BiG-SCAPE at its default cutoff resolves families more finely than both other methods.

**Supplementary Table S4:** GCF inference on curated datasets with expert-assigned families. V-measure at each method's default threshold. $n$: curated BGCs; GCFs: curated families. BiG-SLiCE parsed 261 of the 281 BGCs of MIBiG_BS1.

| Dataset | $n$ | GCFs | This work | BiG-SLiCE | BiG-SCAPE |
|---|---|---|---|---|---|
| Amycolatopsis | 1,200 | 404 | 0.923 | 0.922 | 0.922 |
| Divergent | 39 | 23 | 0.909 | 0.919 | 0.897 |
| JK1 | 279 | 35 | 0.941 | 0.935 | 0.963 |
| MIBiG_BS1 | 281 | 90 | 0.896 | 0.912 | 0.895 |
| Metcalf | 110 | 10 | 0.879 | 0.887 | 0.927 |
| Planomonospora | 11 | 3 | 0.720 | 0.720 | 0.634 |
| Reclassification | 90 | 10 | 0.770 | 0.754 | 0.756 |
| Rhodococcus | 393 | 32 | 0.809 | 0.808 | 0.809 |
| Mean | | | 0.856 | 0.857 | 0.850 |

**Supplementary Table S5:** Runtime of GCF inference. Wall-clock time is the sum over the sequential stages of each run (queue waits excluded); CPU-hours are allocated threads × wall-clock time, summed over all jobs, including parallel array jobs. All runs used 96 threads per job (our procedure: 32–96 threads per stage).

| | This work | | BiG-SLiCE | | BiG-SCAPE | |
|---|---|---|---|---|---|---|
| Input (BGCs) | Wall (h) | CPU-h | Wall (h) | CPU-h | Wall (h) | CPU-h |
| Subset (101,589) | 1.3 | 149 | 1.0 | 97 | 11.3 | 1,080 |
| Subset (507,518) | 2.6 | 670 | 6.3 | 607 | – | – |
| Subset (1,013,058) | 4.0 | 1,322 | 20.3 | 1,952 | – | – |
| antiSMASH-DB 5 (528,998) | 11.5 | 1,146 | 16.1 | 1,542 | >10.7[a] | >1,030 |

[a] Exceeded the 1.9 TB memory limit and was terminated.

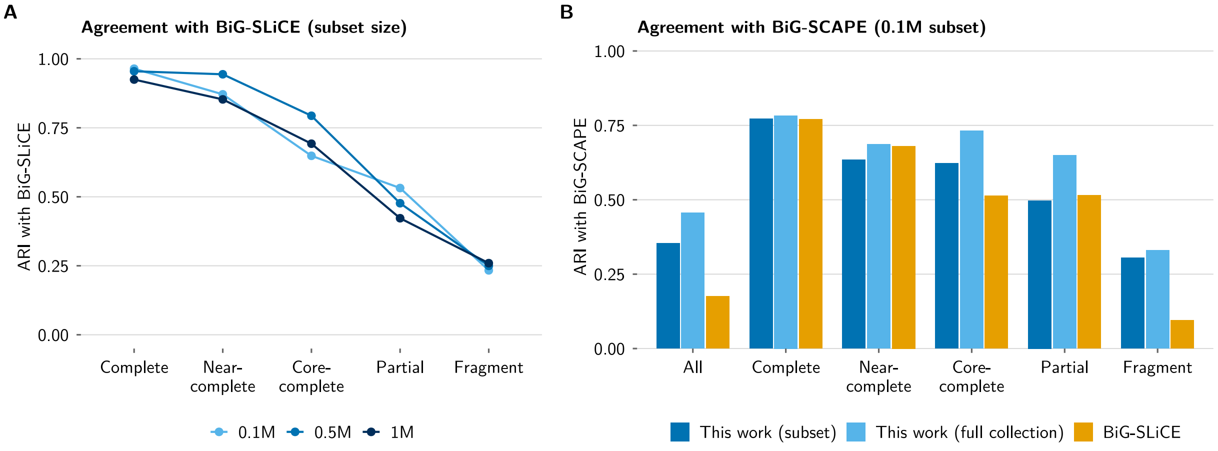


**Supplementary Figure S5:** Agreement of GCF assignments on nested, stratified subsets of BGC Atlas v2. (**A**) Adjusted Rand index (ARI) between this work and BiG-SLiCE 2.0 for the three subset sizes, by length-based completeness class (Supplementary Section S1). (**B**) ARI of this work (run on the subset in isolation, or as assigned in the full collection) and of BiG-SLiCE with BiG-SCAPE 2.0 as reference, on the 0.1-million subset.

**Supplementary Table S6:** Origin of the BGCs in BGC Atlas v2 relative to the first release. First-release assemblies were linked to v2 through their MGnify analysis accessions (33,927 of the corresponding v2 assemblies). BGCs on these assemblies were assigned to contigs that carried at least one first-release BGC or to contigs on which the first release had identified no BGC. Terpene-precursor regions are regions annotated only as terpene-precursor, a region type introduced in antiSMASH 8. The first release contained 1,854,079 BGCs.

| Origin | BGCs | % of v2 |
|---|---|---|
| *Reanalysis of first-release assemblies* | 2,879,693 | 17.9 |
| on contigs with a first-release BGC | 1,613,584 | 10.0 |
| terpene-precursor regions on other contigs | 796,307 | 4.9 |
| other BGCs on other contigs | 469,802 | 2.9 |
| *New data* | 13,229,505 | 82.1 |
| metagenome assemblies | 9,342,160 | 58.0 |
| MAGs | 1,986,083 | 12.3 |
| isolate genomes | 1,900,667 | 11.8 |
| SAGs | 595 | $<0.1$ |
| Total | 16,109,198 | 100.0 |

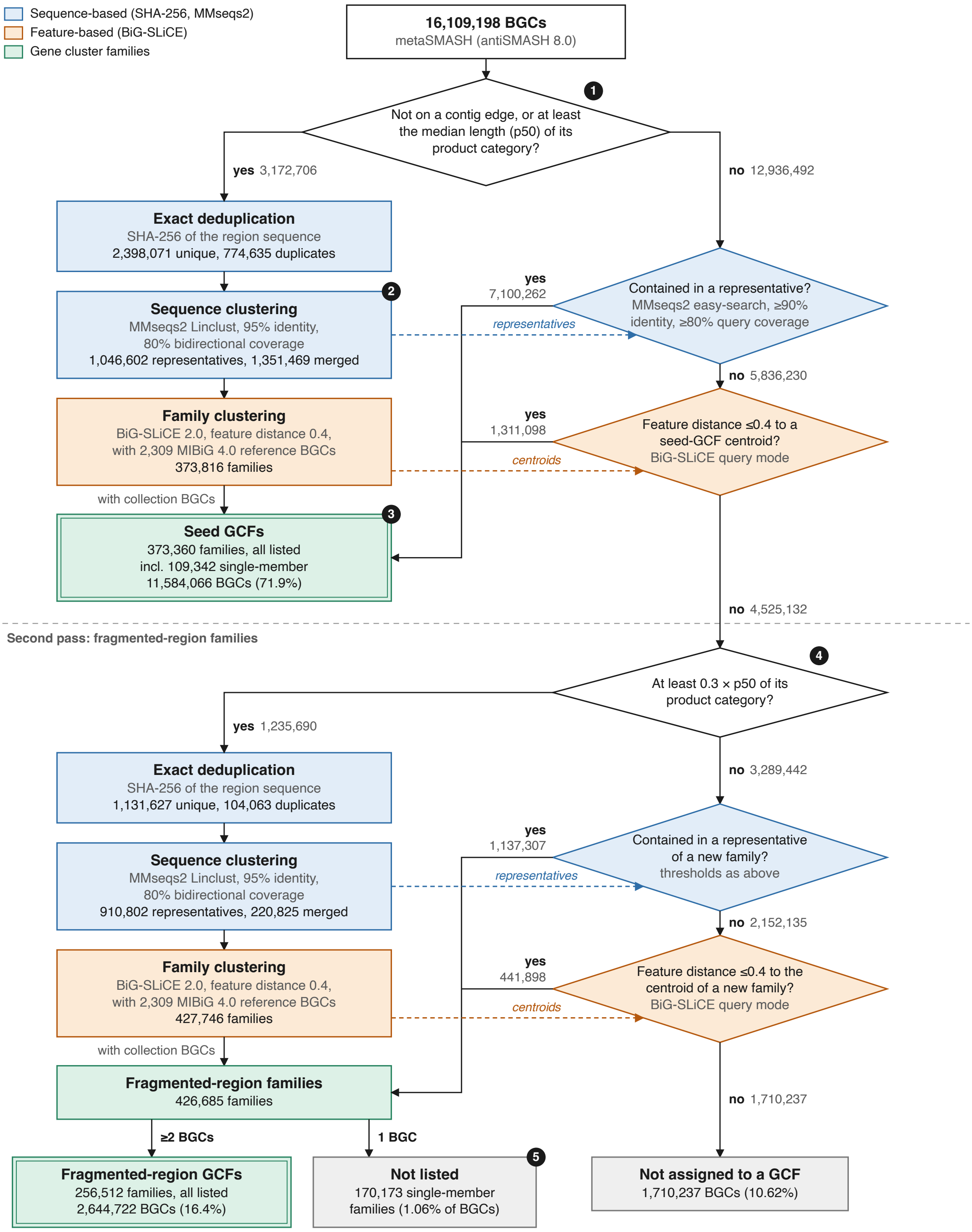


**Supplementary Figure S6:** Gene cluster family (GCF) inference in BGC Atlas v2. BGCs collapsed by deduplication or merged by sequence clustering take the family of their representative. Dashed arrows mark the clustering output that each assignment step searches; grey boxes are BGCs outside listed GCFs, and double borders mark listed GCFs. Arrow counts are from the clustering run and final-box counts from the released catalogue. Design choices: (1) seeds are complete or at least the median length of complete BGCs of their product category (p50; Supplementary Table S3), so they likely retain their biosynthetic core; (2) a sequence cluster's representative is its member not on a contig edge, otherwise its longest member; (3) single-member seed families are listed, as one complete BGC can represent a unique pathway; (4) BGCs shorter than 0.3×p50 do not found families and are only assigned to existing ones; (5) single-member fragmented-region families are not listed, as one fragment is too little evidence for a family. In total, 629,872 listed GCFs contain 14,228,788 BGCs (88.3%).
